# A phylogenetic transform enhances analysis of compositional microbiota data

**DOI:** 10.1101/072413

**Authors:** Justin D Silverman, Alex Washburne, Sayan Mukherjee, Lawrence A David

## Abstract

High-throughput DNA sequencing technologies have revolutionized the study of microbial communities (microbiota) and have revealed their importance in both human health and disease. However, due to technical limitations, data from microbiota surveys reflect the relative abundance of bacterial taxa and not their absolute levels. It is well known that applying common statistical methods, such as correlation or hypothesis testing, to relative abundance data can lead to spurious results. Here, we introduce the PhILR transform, a data transform that utilizes microbial phylogenetic information. This transform enables off-the-shelf statistical tools to be applied to microbiota surveys free from artifacts usually associated with analysis of relative abundance data. Using environmental and human-associated microbial community datasets as benchmarks, we find that the PhILR transform significantly improves the performance of distance-based and machine learning-based statistics, boosting the accuracy of widely used algorithms on reference benchmarks by 90%. Because the PhILR transform relies on bacterial phylogenies, statistics applied in the PhILR coordinate system are also framed within an evolutionary perspective. Regression on PhILR transformed human microbiota data identified evolutionarily neighboring bacterial clades that may have differentiated to adapt to distinct body sites. Variance statistics showed that the degree of covariation of bacterial clades across human body sites tended to increase with phylogenetic relatedness between clades. These findings support the hypothesis that environmental selection, not competition between bacteria, plays a dominant role in structuring human-associated microbial communities.

## INTRODUCTION

Microbiota research today embodies the data-rich nature of modern biology. Advances in high-throughput DNA sequencing allow for rapid and affordable surveys of thousands of bacterial taxa across hundreds of samples (1). The exploding availability of sequencing data has poised microbiota research to advance our understanding of fields as diverse as ecology, evolution, medicine, and agriculture (2). Considerable effort now focuses on interrogating microbiota datasets to identify relationships between bacterial taxa, as well as between microbes and their environment.

Increasingly, it is appreciated that the relative nature of microbial abundance data in microbiota studies can lead to spurious statistical analyses (3-9). With next generation sequencing, the number of reads per sample can vary independently of microbial load (6, 9). In order to make measurements comparable across samples, most studies therefore analyze the relative abundance of bacterial taxa. Analyses are thus not carried out on absolute abundances of community members (**Fig. 1A**), but rather on relative data occupying a constrained, non-orthogonal, geometric space (**Fig. 1B**). Such relative abundance datasets are often termed compositional. The use of most standard statistical tools (*e.g.*, correlation, regression, or classification) within a compositional space leads to spurious results (10). For example, three-quarters of the significant bacterial interactions inferred by Pearson correlation on a compositional human microbiota dataset were likely false (4), and over two-thirds of differentially abundant taxa inferred by a t-test on a simulated compositional human microbiota dataset were spurious (11). To account for compositional effects in microbial datasets, bioinformatics efforts have re-derived common statistical methods including correlation statistics (4, 12), hypothesis testing (13), and variable selection (14, 15).

**Fig. 1.**
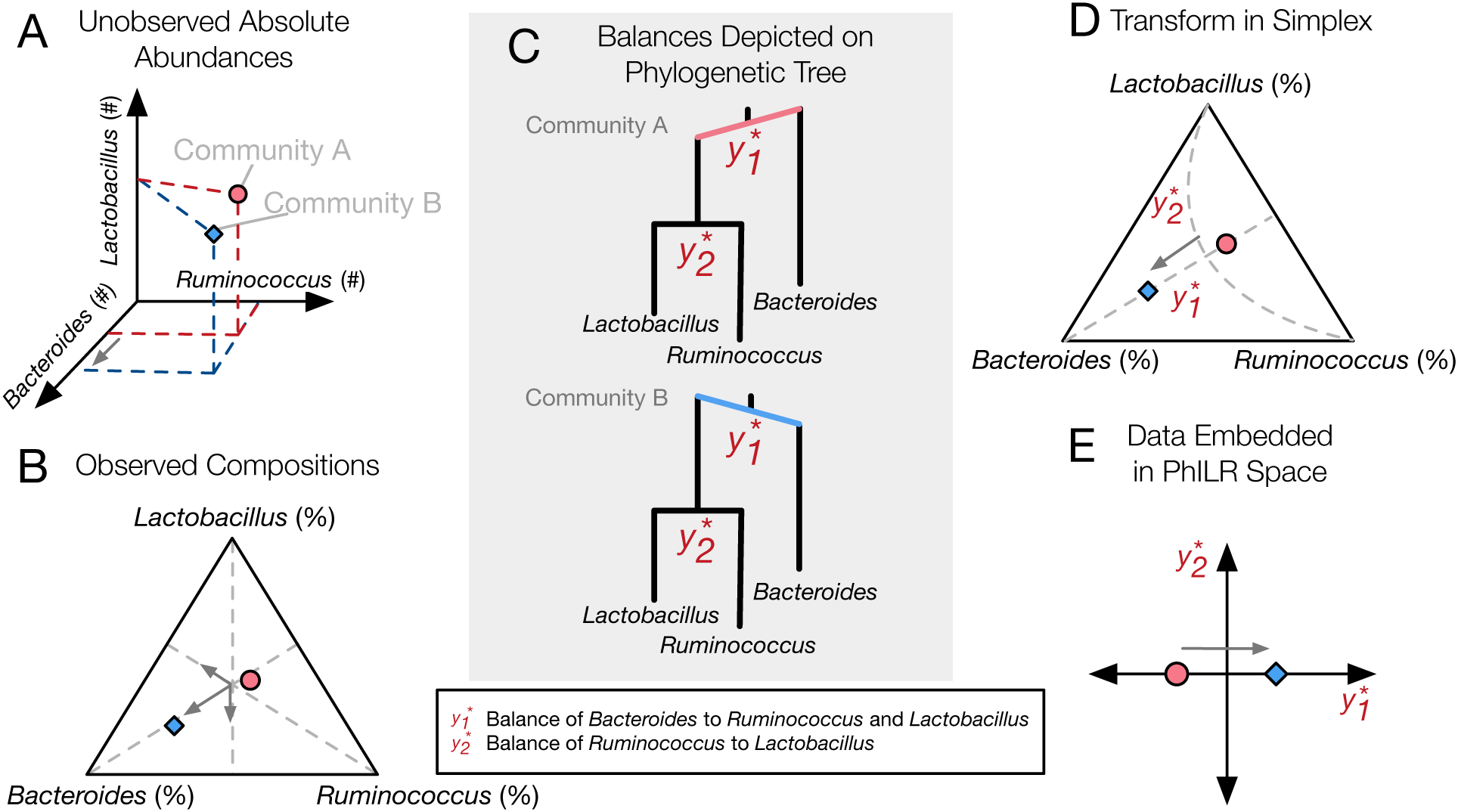
PhILR uses an evolutionary tree to embed microbiota data into an orthogonal, phylogenetically informed space. (**A**) Two hypothetical bacterial communities share identical absolute numbers of *Lactobacillus*, and *Ruminococcus* bacteria; they differ only in the absolute abundance of *Bacteroides* which is higher in community A (red circle) compared to community B (blue diamond). (**B**) A ternary plot depicts proportional data typically analyzed in a sequencing-based microbiota survey. Note that viewed in terms of proportions the space is constrained and the axes are not orthogonal. As a result, all three genera have changed in relative abundance between the two communities. (**C**) Schematic of the PhILR transform based on a phylogenetic sequential binary partition. The PhILR coordinates can be viewed as ‘balances’ between the weights (relative abundances) of the two subclades of a given internal node. In community B, the greater abundance of *Bacteroides* tips the balance 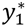 to the right. (**D**) The PhILR transform can be viewed as a new coordinate system (grey dashed lines) in the proportional data space. (**E**) The data transformed to the orthogonal PhILR space. Note that in contrast to the raw proportional data (**B**), the PhILR space only shows a change in the variable associated with *Bacteroides*.

An alternative approach is to transform compositional microbiota data to a space where existing statistical methods may be applied without introducing spurious conclusions. This approach is attractive because of its efficiency: the vast toolbox of existing statistical models can be applied without re-derivation. Normalization methods, for example, have been proposed to modify count data by assuming reads follow certain statistical distributions (*e.g.*, negative binomial) (16, 17). Alternatively, the field of Compositional Data Analysis (CoDA) has focused on formalizing methods for transforming compositional data from a constrained non-orthogonal space into a simpler geometry without having to assume data adhere to a distribution model (18). Previous microbiota analyses have already leveraged CoDA theory and used the centered log-ratio transform to reconstruct microbial association networks and interactions (19, 20) and to analyze differential abundances (21, 22). However, the centered log-ratio transform has a crucial limitation: it yields a coordinate system featuring a singular covariance matrix and is thus unsuitable for many common statistical models (10). This drawback can be sidestepped using another CoDA transform, known as the Isometric Log-Ratio (ILR) transformation (23). The ILR transform uses a sequential binary partition of the original variable space (**Fig. 1C**) to create a new coordinate system with orthonormal bases (**Fig. 1D,E**) (23). However, a known obstacle to using the ILR transform is the choice of partition such that the resulting coordinates are meaningful (10). To date, microbiota studies have chosen ILR coordinates using random sequential binary partitions of bacterial groups (24, 25).

Here, we introduce the bacterial phylogenetic tree as a natural and informative sequential binary partition when applying the ILR transform to microbiota datasets (**Fig. 1C**). Using phylogenies to construct the ILR transform results in an ILR coordinate system capturing evolutionary relationships between neighboring bacterial groups (clades). Analyses of neighboring clades offer the opportunity for biological insight: clade analyses have linked genetic adaptation to ecological differentiation (26), and the relative levels of sister bacterial genera differentiate human cohorts by diet, geography, and culture (27-29). Datasets analyzed by a phylogenetically aware ILR transform could therefore reveal ecological and evolutionary factors shaping host-associated microbial communities.

We term our approach the **Ph**ylogenetic **ILR** (PhILR) transform. Using environmental and human-associated 16S rRNA studies as benchmarks, we show that the accuracy of distance-based and machine learning models often increases and never decreases after applying the PhILR transform, relative to applying the same models on untransformed (raw) or log transformed relative abundance data. Moreover, because the PhILR transform incorporates phylogenetic information, statistics applied in the PhILR coordinate system naturally identify bacterial clades that may have differentiated to adapt to distinct body sites. The PhILR coordinate system can also be used to show that, in all human body sites studied, the degree to which neighboring bacterial clades covary tends to increase with the phylogenetic relatedness between clades. This result supports theories that environmental forces, and not competition between bacteria, primarily structure the assembly of human microbiota.

## RESULTS

### Constructing the PhILR transform

The PhILR transform has two goals. The first goal is to transform input microbiota data into an unconstrained orthogonal space while preserving all information contained in the original composition. The second goal is to conduct this transform using phylogenetic information. To achieve these dual goals on a given set of *N* samples consisting of relative measurements of *D* taxa (**Fig. 1B**), we transform data into a new space of *N* samples and (*D* − 1) coordinates termed ‘balances’ (**Fig. 1C-E**). Each balance 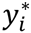 is associated with a single internal node *i* of a phylogenetic tree with the *D* taxa as leaves. The balance represents the log-ratio of the geometric mean relative abundance of the two clades of taxa that descend from *i* (see *Methods* and *SI Text*). Balances are by definition orthogonal, which ensures standard statistical tools may be applied to the transformed data without compositional artifacts; however, this orthogonality does not imply statistical independence (*SI Text*). Each balance is also standardized so that balances across the tree are statistically comparable (10), even when balances have differing numbers of descendant tips or exist at different depths in the tree. This standardization also ensures that the variance of PhILR balances has a consistent scale, unlike the variance of standard log-ratios where it is often unclear what constitutes a large or small variance (4).

### Benchmarking statistics in the PhILR coordinate system

To assess how the PhILR transform affects statistical inference on microbiota datasets, we first examined measures of community dissimilarity. Microbiota analyses commonly compute the dissimilarity or distance between pairs of samples and identify groups of samples with differing community structure. We benchmarked how the PhILR transform affected the task of grouping samples using three microbiota surveys as references: Costello Skin Sites (CSS), a dataset of 357 samples from 12 human skin sites (30); Human Microbiome Project (HMP), a dataset of 4,743 samples from 18 human body sites (*e.g*., skin, vaginal, oral, and stool) (31); and, Global Patterns (GP), a dataset of 26 samples from 9 human or environmental sites (1) (**Fig. S1**). We computed distances between samples in the PhILR coordinate system using Euclidean distances. We compared this measure to common measures of microbiota distance or dissimilarity (Unifrac, Bray-Curtis and Jaccard) as well as a simple measure (Euclidean) applied to raw relative abundance data (32).

The PhILR transform significantly improved distance-based analyses of microbiota samples. Principal coordinate analyses (PCoA) qualitatively demonstrated separation of body sites using both Euclidean distances on PhILR transformed data (**Fig. 2A**) and with a number of standard distance measures calculated on raw relative abundance data (**Fig. S2**). To quantitatively compare distance measures, we tested how well habitat information explained variability among distance matrices using PERMANOVA (33). The Euclidean distance in the PhILR coordinate system significantly outperformed the five competing distance metrics across all benchmarks, except in comparison to Weighted Unifrac when applied to the HMP dataset (**Fig. 2B**). These results indicate that the Euclidean distance, when measured in the PhILR coordinate system, generally exhibited superior performance to more sophisticated distance measures used on raw relative abundances.

**Fig. 2.**
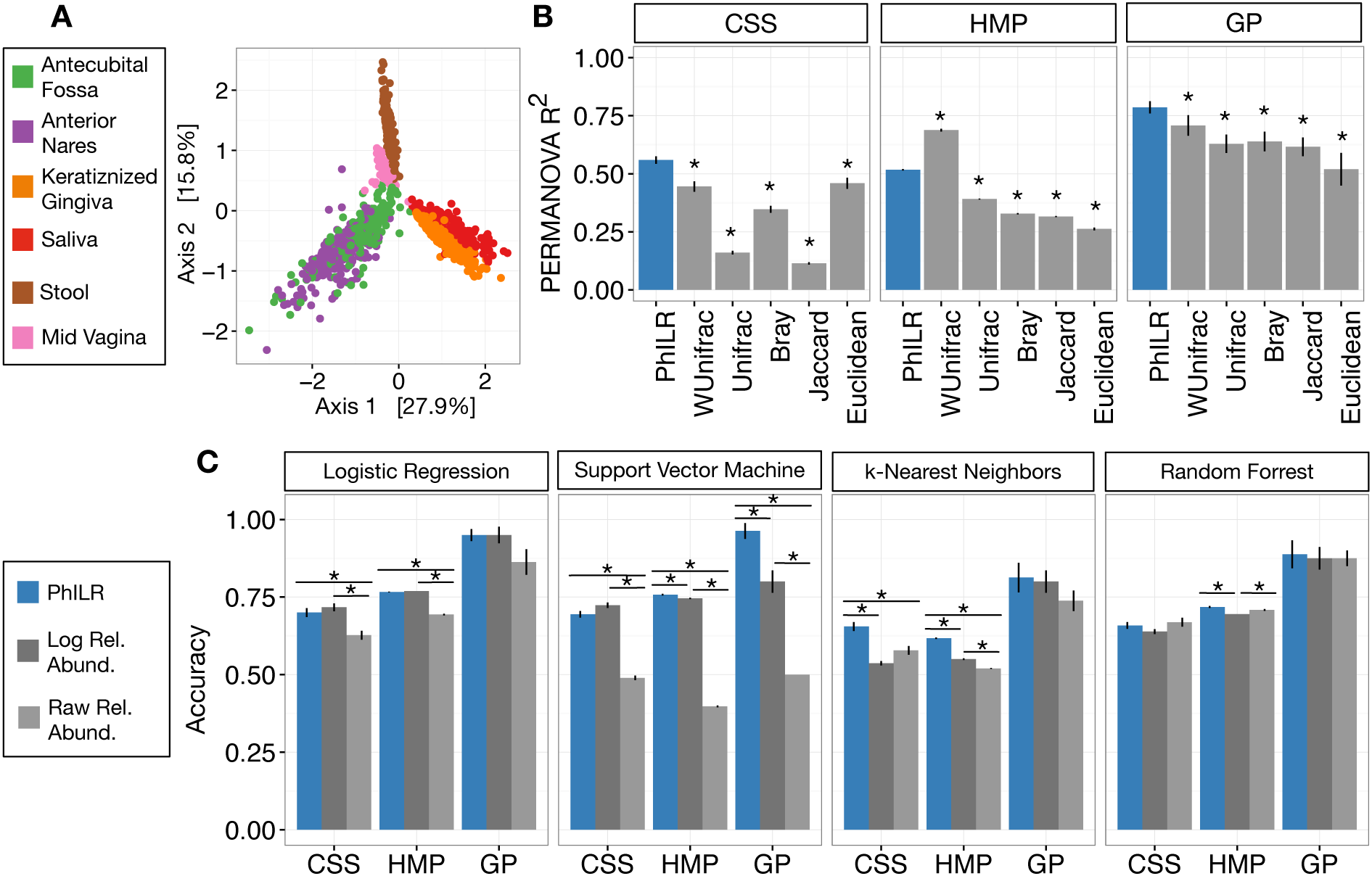
PhILR transform improves performance of standard statistical models on microbiota data. Benchmarks were performed using three datasets: Costello Skin Sites(CSS), Global Patterns (GP), Human Microbiome Project (HMP). (**A**) Sample distance visualized using principal coordinate analysis (PCoA) of Euclidean distances computed in PhILR coordinate system. A comparison to PCoAs calculated with other distance measures is shown in **Fig. S2**. (**B**) Sample distance (or dissimilarity) was computed by a range of statistics. R^2^ values from PERMANOVA were used to measure how well sample location explained the variability in distances between samples. Distances in the PhILR transformed space were calculated using Euclidean distance. Distances between samples on raw relative abundance data were computed using Weighted and Unweighted UniFrac (WUnifrac and Unifrac, respectively), Bray-Curtis, Binary Jaccard, and Euclidean distance. Error bars represent standard error measurements from 100 bootstrap replicates and (*) denotes a p-value of ≤0.01 after FDR correction of pairwise tests against PhILR. (**C**) Accuracy of supervised classification methods tested on benchmark datasets. The PhILR transform significantly improved supervised learning algorithm accuracy in 7 out of 12 supervised classification benchmarks compared to raw or log-transformed raw data. Error bars represent standard error measurements from 10 test/train splits and (*) denotes a p-value of !0.01 after FDR correction of all pairwise tests.

Next, we tested the performance of predictive statistical models in the PhILR coordinate system. We examined four standard supervised machine learning techniques: logistic regression (LR), support vector machines (SVM), k-nearest neighbors (kNN), and random forests (RF) (34). We applied these methods to the same three reference datasets used in our comparison of distance metrics. As a baseline, the machine learning methods were applied to raw relative abundance datasets and raw relative abundance data that had been log-transformed.

The PhILR transform significantly improved supervised classification accuracy in 7 of the 12 benchmark tasks compared to raw relative abundances (**Fig. 2C**). Accuracy improved by more than 90% in two benchmarks (SVM on HMP and GP), relative to results on the raw data. Log transformation of the data also improved classifier performance significantly on 6 of the 12 benchmarks but also significantly underperformed on 1 benchmark compared to raw relative abundances. In addition, the PhILR transform significantly improved classification accuracy in 5 of the 12 benchmarks relative to the log transform. Overall, the PhILR transform often outperformed the raw and log transformed relative abundances with respect to classification accuracy and was never significantly worse.

### Identifying neighboring clades that differ by body site preference

We next used a sparse logistic regression model to examine which balances distinguished human body site microbiota in the HMP dataset. Such balances could be used to identify neighboring bacterial clades whose relative abundances capture community-level differences between body site microbiota. Microbial genetic differentiation may be driven by adaptation to new resources or lifestyle preferences (26), meaning that distinguishing balances near the tips of the bacterial tree may correspond to clades adapting to human body site environments.

We identified dozens of highly discriminatory balances, which were spread across the bacterial phylogeny (**Fig. 3A** and **Fig. S3, S4**). Some discriminatory balances were found deep in the tree. Abundances of the Firmicutes, Bacteroidetes, and Proteobacteria relative to the Actinobacteria, Fusobacteria, and members of other phyla, separated skin body sites from oral and stool sites (**Fig. 3B**). Levels of the genus *Bacteroides* relative to the genus *Prevotella* differentiated stool microbiota from other communities on the body (**Fig. 3C**). Notably, values of select balances below the genus level also varied by body site. Relative levels of sister *Corynebacterium* species separated human skin sites from gingival sites (**Fig. 3D**). Species-level balances even differentiated sites in nearby habitats; levels of sister *Streptococcus* species or sister *Actinomyces* species vary depending on specific oral sites (**Fig. 3E,F**). These results show that the PhILR transform can be used to highlight ancestral balances that distinguish body site microbiota, as well as to identify more recent balances that may separate species that have adapted to inhabit different body sites.

**Fig. 3.**
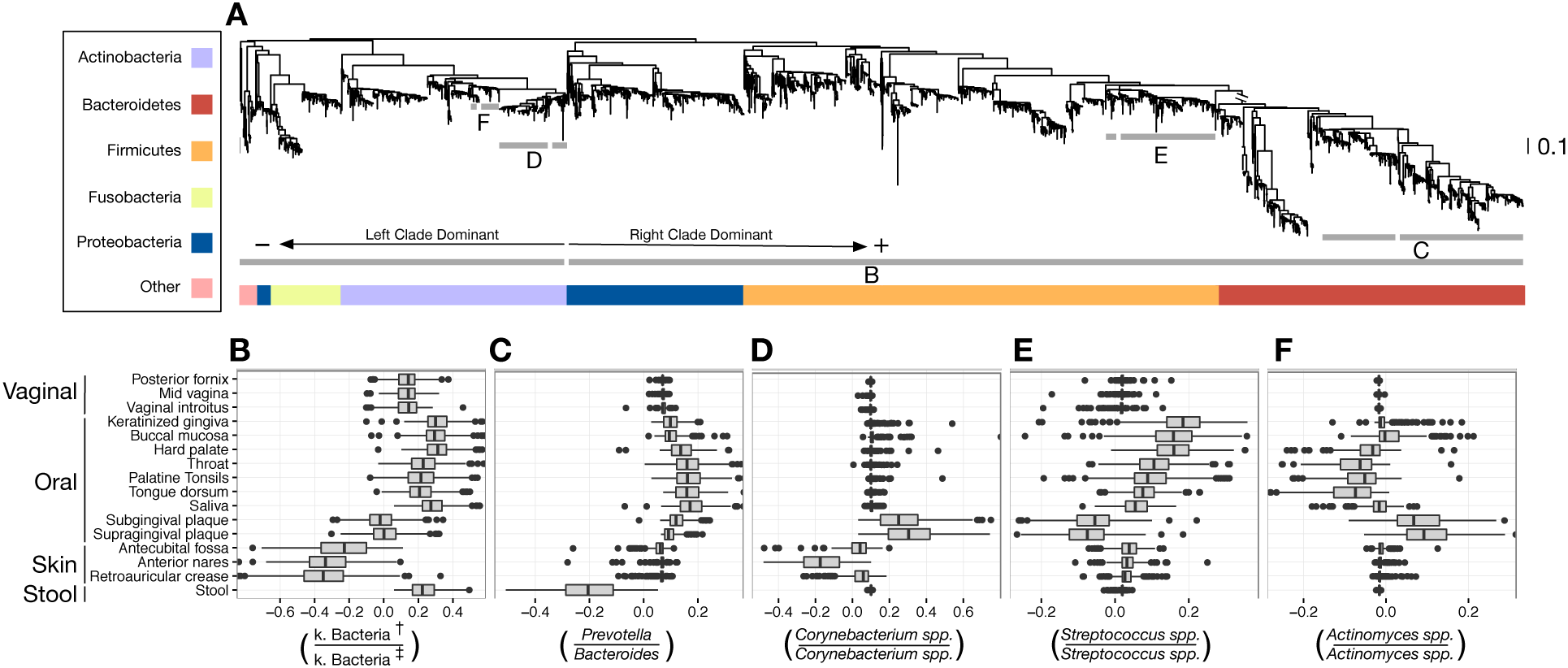
Balances distinguishing human microbiota by body site. Sparse logistic regression was used to identify balances that best separated the different sampling sites (full list of balances provided in **Fig. S3-4**). (**A**) Each balance is represented on the tree as a broken grey bar. The left portion of the bar identifies the clade in the denominator of the log-ratio, and the right portion identifies the clade in the numerator of the log-ratio. The branch leading from the Firmicutes to the Bacteroidetes has been rescaled to facilitate visualization. (**B**-**F**) The distribution of balance values across body sites. Vertical lines indicate median values, boxes represent interquartile ranges (IQR) and whiskers extend to 1.5 IQR on either side of the median. Balances between: (**B**) the phyla Actinobacteria and Fusobacteria versus the phyla Bacteroidetes, Firmicutes, and Proteobacteria distinguish stool and oral sites from skin sites; (**C**) *Prevotella spp.* and *Bacteroides spp.* distinguish stool from oral sites; (**D**) *Corynebacterium spp.* distinguish skin and oral sites; (**E**) *Streptococcus spp*. distinguish oral sites; and, (**F**) *Actinomyces spp.* distinguish oral plaques from other oral sites. (†) Includes Bacteroidetes, Firmicutes, Alpha-, Beta-, and Gamma-proteobacteria. (‡) Includes Actinobacteria, Fusobacteria, Epsilon-proteobacteria, Spirochaetes, and Verrucomicrobia.

### Balance variance and microbiota assembly

Observing that discriminatory balances could be found across the phylogenetic tree suggested investigating theories of microbial community assembly within the PhILR coordinate system. Closely related microbes may directly compete for nutrients and thus exclude one another from a given site (35). By contrast, related taxa may also have similar lifestyle characteristics and thus covary in environments favoring their shared traits (36). Patterns of phylogenetic clustering (36) or predicted metabolic interactions (37, 38) have previously been used to distinguish the relative importance of competition and environmental selection in structuring microbial communities (36).

The variance of a balance in the PhILR coordinate system provides an alternative phylogenetic method to measure how bacterial taxa covary across environments. In contrast to standard measures of association (*e.g*., Pearson correlation), balance variance is robust to compositional artifacts (10). When the variance of a balance between two clades approaches zero, the mean abundance of taxa in each of the two clades will be linearly related and thus exhibit shared dynamics across microbial habitats (39). By contrast, when a balance exhibits high variance, related bacterial clades exhibit unlinked or exclusionary dynamics across samples. A pattern of lower balance variance near the tips of the phylogenetic tree would suggest that closely related taxa tend to covary and support the hypothesis that environmental forces structure sampled microbial communities; by contrast, higher balance variance near the tips of the phylogeny would suggest related taxa do not covary and support a competitive model underlying community structure.

For all body sites in the HMP dataset, we observed significantly decreasing balance variances near the tips of the phylogenetic tree (p<0.01, permutation test with FDR correction; *Methods*; **Fig. 4A-F** and **Fig. S5, S6**). Low variance balances predominated near the leaves of the tree. Examples of such balances involved *B. fragilis* species in stool (**Fig. 4H**), *Rothia mucilaginosa* species in the buccal mucosa (**Fig. 4J**), and *Lactobacillus* species in the mid-vagina (**Fig. 4L**). By contrast, higher variance balances tended to be more basal on the tree. Two examples of high variance balances corresponded with clades at the order (**Fig. 4G**) and family (**Fig. 4I**) levels. In the case of select Lactobacilli in the vagina, neighboring clades appeared to exclude one another (**Fig. 4K**). We performed LOESS regression to investigate how the relationship between balance variance and phylogenetic depth varied locally at different taxonomic scales. This regression revealed that trends between variance and phylogenetic depth were stronger above the species level than below this level (*Methods*; **Fig. 4D-F** and **Fig. S6**). Overall, the observed pattern of decreasing balance variance near the tips of the phylogenetic tree demonstrated that closely related bacteria tend to covary in human body sites, supporting the hypothesis that environmental forces structure human-associated microbial communities more than competitive forces. However, the weaker relationship between balance variance and phylogenetic depth below the species level suggests that environmental forces induce similar selective pressure on bacterial strains within the same species.

**Fig. 4.**
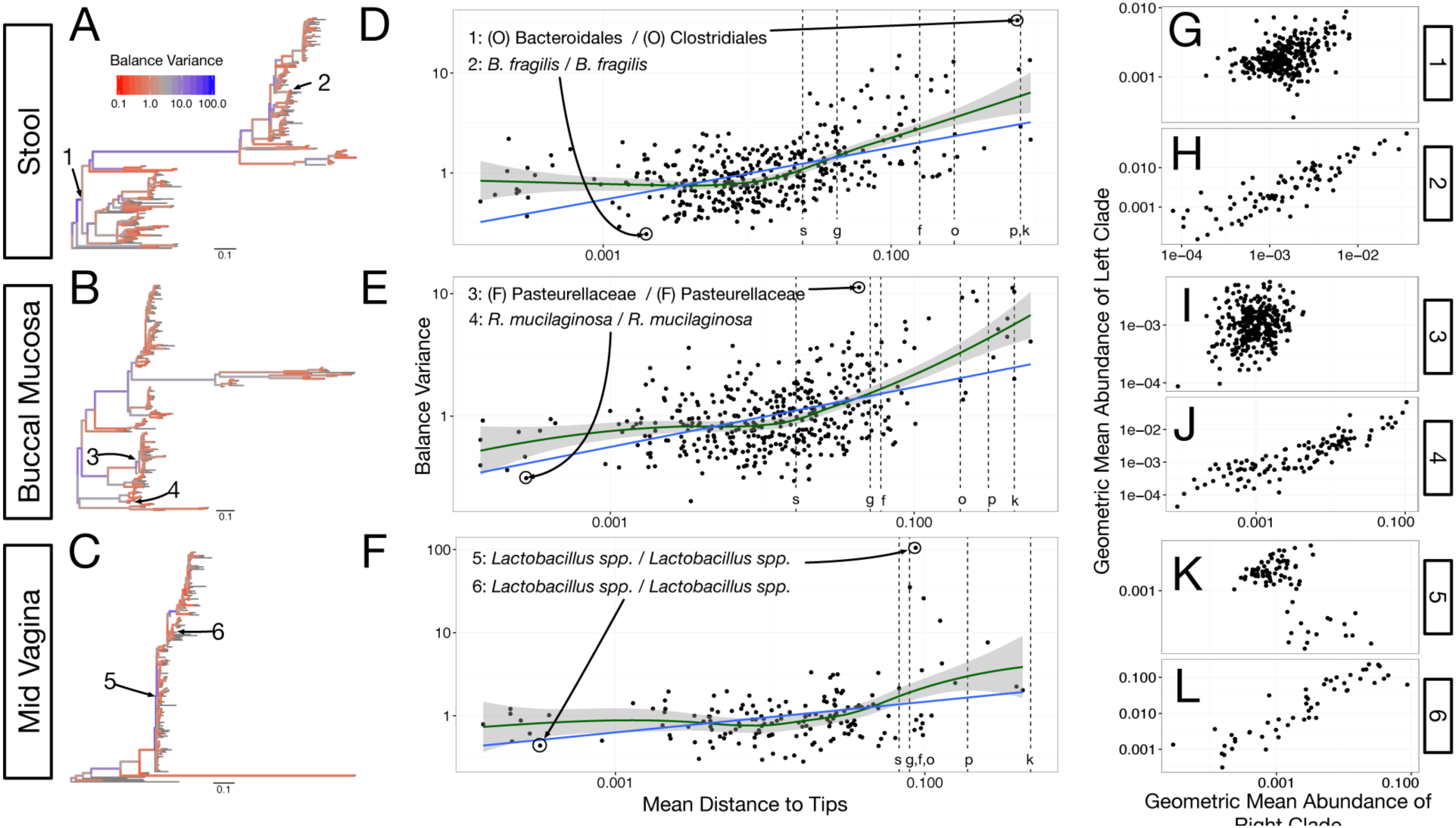
Neighboring clades covary less with increasing phylogenetic depth. The variance of balance values captures the degree to which neighboring clades covary, with smaller balance variances representing sister clades that covary more strongly. (**A**-**C**) Balance variances were computed among samples from stool (**A**), buccal mucosa (**B**), and the mid-vagina (**C**). Red branches indicate small balance variance and blue branches indicate high balance variance. Balances 1-6 are individually tracked in panels (**D-L**). (**D**-**F**) Balance variances within each body site increased linearly with increasing phylogenetic depth on a log-scale (blue line; p<0.01, permutation test with FDR correction). Significant trends are seen across all other body sites (**Fig. S6**). Non-parametric LOESS regression (green line and corresponding 95% confidence interval) reveals an inflection point in the relation between phylogenetic depth and balance variance. This inflection point appears below the estimated species level (‘s’ dotted line; the median depth beyond which balances no longer involve leaves sharing the same species assignment; *Methods*). (**G**-**L**) Examples of balances with high and low variance from panels (**A**-**F**). Low balance variances (**H, J, L**) reflect a linear relationship between the geometric means of sister clades abundances. High balance variances reflect either unlinked (**G, I**) or exclusionary (**K**) dynamics between the geometric means of sister clades abundances.

## DISCUSSION

The relative nature of microbiota survey data can result in spurious statistical analyses. Here, we addressed this problem by developing a technique to transform conventional microbiota data into a new space free from compositional effects. The resulting data space improves the accuracy of common statistical methods when applied to microbiota data. The PhILR transform also embeds phylogenetic information into statistics computed in its coordinate system. In doing so, the PhILR transform provides a natural means for discovering taxonomic and evolutionary factors structuring microbial communities.

Our benchmarking experiments show that relative to untransformed data, the PhILR transform improves the performance of both distance-based and supervised machine learning algorithms applied to microbiota data. We note that performance gains achieved by the PhILR transform on community distance benchmarks are surpassed in only one instance by the phylogenetic distance Weighted Unifrac. Unifrac down weights the influence of closely related taxa when computing the distance between communities, and a similar effect can be achieved when Euclidean distances are calculated with PhILR transformed data (*Methods*). Because related bacteria often share similar traits (40), this weighting likely biases the grouping of microbiota so that communities with similar functional profiles are nearby in the transformed space. Our benchmarking suggests that in practice, leveraging phylogenetic information and accounting for compositional constraints can improve statistical analysis of microbiota surveys.

The PhILR transform’s use of phylogenetic information also helps formalize the practice of distinguishing microbiota using taxonomic ratios. For example, the enteric Firmicutes to Bacteroidetes ratio has repeatedly been compared between obese and lean individuals (41-44). In part, such ratios are relied on because they simplify microbiota with hundreds of component species into single variables. The PhILR transform provides a statistical framework that guides the process of finding pairs of bacterial clades that differentiate groups of samples. Important balances identified correspond to ratios already known to distinguish microbiota in practice; *e.g*. relative abundances of Actinobacteria to other bacterial phyla, which we find separate skin samples from other human body sites (**Fig. 3B**), have previously been used to identify skin microbiota (45). Interestingly, the PhILR transform also identifies new uses for well-known ratios. The balance between the genera *Bacteroides* and *Prevotella*, which has been previously linked to inter-individual stool variation (46, 47), emerged as one of the best discriminants separating human stool samples from other body sites (**Fig. 3C**). This finding was likely sensitive to the use of a Western subject cohort in the Human Microbiome Project; a cohort drawn from non-industrialized settings would likely have exhibited higher levels of enteric *Prevotella* (27). Nevertheless, we anticipate the PhILR coordinate system to be a useful tool for identifying clades of bacteria that vary by habitat.

Although discriminatory balances between habitats could be constructed between unrelated clades, the PhILR transform’s reliance on phylogenetically defined balances also carries the benefit of linking subsequent statistical analyses to evolutionary models. A symbiosis exists between our understanding of bacterial evolution and the ecology of host-associated microbial communities (48). Microbiota studies have shown that mammals and bacteria cospeciated over millions of years (49, 50), and human gut microbes have revealed the forces driving horizontal gene transfer between bacteria (51). Evolutionary tools and theory have been used to explain how cooperation benefits members of gut microbial communities (52), and raise concerns that rising rates of chronic disease are linked to microbiota disruption (53). The PhILR transform provides a convenient framework for carrying out statistical analyses in a coordinate system that is evolutionarily informed.

Regression on PhILR transformed data, for example, highlighted balances near the tips of the bacterial phylogeny that distinguished human body sites. These balances may reflect functional specialization, as ecological partitioning among recently differentiated bacterial clades could be caused by genetic adaptation to new environments or lifestyles (26). Indeed, among oral body sites, we observed consistent site specificity of neighboring bacterial clades within the genera *Actinomyces* (**Fig. 3F**) and *Streptococcus* (**Fig. 3E**). Species within the *Actinomyces* genera have been previously observed to partition separately between the teeth, gingival plaque, buccal mucosa and tongue in healthy subjects (54, 55). Even more heterogeneity has been observed within the *Streptococcus* genus, where species have been identified that distinguish teeth, plaque, mucosal, tongue, saliva, and other oral sites (54, 55). This partitioning likely reflects variation in the anatomy and resource availability across regions of the mouth (54), as well as the kinds of surfaces bacterial strains can adhere to (55).

We also observed evidence for potential within-genus adaptation to body sites that has not been previously reported. In particular, within the genus *Corynebacterium*, we found ratios of taxa varied among oral plaques and select skin sites (**Fig. 3D**). Although the genus is now appreciated as favoring moist skin environments, the roles played by individual Corynebacteria within skin microbiota remain incompletely understood (45). Precisely linking individual *Corynebacterium* species or strains to body sites is beyond the scope of this study due to the limited taxonomic resolution of 16S rRNA datasets (56, 57). Nevertheless, we believe the PhILR coordinate system may be used in the future to identify groups of related bacterial taxa that have undergone recent functional adaptation.

Another example of how the PhILR transform may be used to provide evolutionary insight arises in our analysis of balance variance and phylogenetic depth. The relative importance of environmental and competitive forces in shaping human microbiota remains an outstanding question for microbial ecology (35). Reports of paired strains within the same genus or species that inhibit growth of one another (58-62) have suggested that competitive forces are dominant. By contrast, we observed decreasing balance variance near the tips of the phylogenetic tree (**Fig. 4A-F** and **Fig. S6**), supporting the hypothesis that microbiota in different body sites are shaped primarily by environmental forces. Such forces could include moisture, oxygen level, or resource availability (45, 63, 64). Our findings complement previous studies that used metabolic interactions to show that in the human gastrointestinal and oral microbiomes, species tend to co-occur with other species with which they strongly compete (37, 38). In addition, our conclusions are supported by recent fecal transplant experiments in humans showing that the presence of conspecific bacteria increases the likelihood that a bacterial strain engrafts in the human gut (65).

We also found that the relationship between balance variance and phylogenetic depth varies with taxonomic scale, appearing stronger at balances corresponding to higher taxonomic scales and weaker at balances near or below the species level. Although it is often believed that microbial phenotypes are linked to phylogenetic distance (40, 66), the precise taxonomic levels at which this relationship degrades remains debated (67). Our phylogenetic analysis suggests that lifestyle characteristics enabling bacteria to persist in human body sites are conserved among strains roughly corresponding to the same species.

Ultimately, though the methods presented here provide a coherent geometric framework for performing microbiota analysis in a compositionally robust manner, future refinements and modifications are possible. As it is often unclear as to when a zero value represents a value below the detection limit (rounded zero) or a truly absent taxa (essential zero), the handling of zero values remains an outstanding challenge for microbiota analysis and compositional data analysis. Here, we have used two methods for handling zeros depending on the biological question of interest (*Methods, SI Text*); however, new mixture models that explicitly allow for both essential and rounded zeros (68) appear promising for microbiota data analysis. Additionally, we chose to use phylogenies to create the sequential binary partitions needed for the ILR transform. This algorithm design choice provided our analyses with evolutionary context, but such context may not be needed for every analysis. Alternative balances between non-phylogenetically neighboring groups of taxa can be constructed and used with the ILR transform, provided the overall partition of the taxa is binary. Lastly, if analytical insights are desired on the level of individual taxa, and not ratios of clades, analysis can be performed in the transformed ILR space and the results then converted back into compositional space using the inverse of the ILR transform (10, 69). This provides an alternative approach to adapting statistical methods for use with compositional microbiota data.

Yet, despite these avenues for improvement or modification, we believe the PhILR transform already enables existing statistical methods to be applied to metagenomic datasets, free from compositional artifacts and framed according to an evolutionary perspective. We emphasize that all statistical tools applied to PhILR transformed data in this study were used 'off-the-shelf' and without modification. This approach contrasts with the more standard practice of adapting current statistical techniques to the limitations of microbiota survey data. Such adaptation is often challenging because many statistics were derived assuming unconstrained orthogonal coordinate systems, not constrained and over-determined compositional spaces. Therefore, while select techniques have already been adapted (*e.g*. distance measures that incorporate phylogenetic information (70) and feature selection methods that handle compositional input (14, 15)), it is likely that certain statistical goals, such as non-linear community forecasting or control system modeling, may prove too complex for adapting to microbiota datasets. Beyond microbiota surveys, we also recognize that compositional metagenomics datasets are generated when studying the ecology of viral communities (71) or clonal population structure in cancer (72-74). We expect the PhILR transform to aid other arenas of biological research where variables are measured by relative abundance and related by an evolutionary tree.

## METHODS

### Overview of the PhILR Transform

The PhILR transform is an Isometric Log-Ratio transform (23) defined by using a binary phylogenetic tree as a sequential binary partition. The PhILR transform also involves an optional scaling step to integrate phylogenetic distances into the transformed space (branch length weighting) and two methods for handling zero values (taxa weighting and conditioning on non-zero counts). We describe these methods and their motivation in more detail below. A more detailed description of their derivation and the underlying theory of the transform is provided in the *SI Text*.

### The ILR Transform

A typical microbiome sample consists of measured counts *c_j_* for taxa *j* ∈ {1,…, *D*}. A standard operation is to take count data and transform it to relative abundances. This operation is referred to as closure in compositional data analysis and is given by

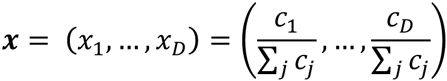

where *x_j_* represents the relative abundance of taxa *j* in the sample. We can represent a binary phylogenetic tree of the *D* taxa using a sign matrix Θ. Each row of the sign matrix indexes an internal node *i* of the tree and each column indexes a tip of the tree. A given element in the sign matrix is ±1 depending on whether that tip is in the left or right subtree descending from *i* and 0 if that tip is not a descendent of *i* (**Fig. S7**). Following Egozcue and Pawlowsky-Glahn (69), we represent the ILR coordinate (balance) associated with node *i* in terms of the shifted composition ***y*** = ***x/p*** = (*x*_1_/*p*_1_,…, *x*_*D*_/*p*_*D*_) as

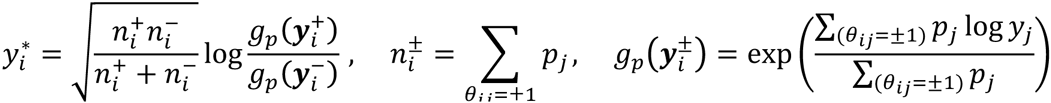

where 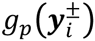 represents the weighted geometric mean of the components of ***y*** that are descendants of the left or right subtree of node *i* respectively and *p*_*j*_ is given by the weights assigned to taxa *j*. When ***p*** = (1,…,1), ***y*** = ***x*** and the above equation represents the ILR transform as originally published (75). However, when ***p*** ≠ (1,…,1), the above equation represents a more generalized form of the ILR transform (69) that allows the effects of very low abundance taxa to be down weighted. *(See the SI Text for more background on the form of this transformation)*

### Addressing sparsity through weighting taxa

We make use of this generalized ILR transform and weights ***p*** to address the challenge of zero and near-zero counts. In most analyses involving count data the challenge of modeling zero or near-zero counts must be addressed. After replacing zeros with small non-zero counts, a standard procedure to address this challenge is variance stabilization, down-weighting the influence of small counts since these are less reliable and therefore more variable (76). Microbiota datasets are often sparse, with more than than 90% zero counts. Through the use of the generalized ILR we may down weight the less reliable rare and very low abundance taxa by adjusting the weights ***p***. This reduces the sensitivity of statistics in the PhILR coordinate system to very low abundance taxa. As total read counts contain variance information (77), we utilize total read counts to choose these weights. We refer to this method as ‘taxa weighting’.

Our choice of taxa weights combines two terms multiplicatively, a mean or median of the raw counts for a taxon across the *N* samples in a dataset and the norm of the vector of relative abundances of a taxon across the *N* samples in a dataset. This second term ensures that highly site-specific taxa are not unduly down weighted (*SI Text*). Preliminary studies showed that the geometric mean of the counts (with a pseudocount added to avoid skew from zero values) outperformed both the arithmetic mean and median as a measure of central tendency for the counts (data not shown). Both the Euclidean norm and the Aitchison norm improved performance (as measured by classification accuracy or PERMANOVA R^2^, see below) in our benchmark tasks as compared to using the geometric mean alone (**Fig. S8**). However in one case (classification using support vector machine on the global patterns dataset) the Euclidean norm greatly outperformed the Aitchison norm and was therefore chosen for our analysis here. Thus, the taxa weights we used are given by

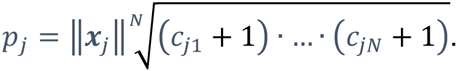

Note that we add the subscript *j* to the right hand side of the above equation to emphasize that this is calculated with respect to a single taxon across the *N* samples in a dataset. These taxa weights supplement the use of pseudo-counts with variance information from total count data and, with the exception of our analysis of balance variance as a function of phylogenetic depth (see below), are used throughout the analyses presented here.

### Incorporating branch lengths

Beyond utilizing the connectivity of the phylogenetic tree to dictate the partitioning scheme for ILR balances, branch length information can be embedded into the transformed space by linearly scaling ILR balances 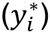 by the distance between neighboring clades. We call this scaling by phylogenetic distance ‘branch length weighting’. This has the effect of scaling distances in the PhILR coordinate system by the relatedness of bacteria present in a community, which is a feature shared by other phylogenetic methods that utilize phylogenetically informed distances (70, 78, 79). However, importantly, whereas those methods are based on reducing the data to a set of distances, the PhILR transform provides an explicit coordinate system of balances where each balance identifies a distinct location on the phylogenetic tree and has evolutionary meaning. Specifically for each coordinate 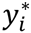, corresponding to node *i* we use the transform

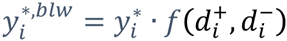

where 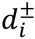 represent the branch lengths of the two direct children of node *i*. When 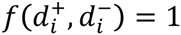, the coordinates are not weighted by branch lengths. With the exception of our analysis of balance variance as a function of phylogenetic depth (see below), here we have used 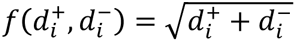 for our branch length weights. When coupled with the taxa weights specified above, the square root of the summed distances had the highest rank in 9 of the 12 supervised classification tasks and 2 of the 3 distance based tasks (compared to either 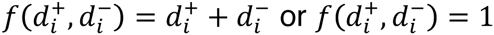; **Fig. S8**). This choice of branch length weights most similarly resembles the generalized UniFrac distance with *d* = 0.5 (33).

### Implementation

The PhILR transform, as well as the incorporation of branch length and taxa weightings has been implemented in the R programing language as the package *philr* available at https://github.com/jsilve24/philr. The implementation is limited in time and space complexity by a single matrix multiplication step involving a *D*×(*D* − 1) contrast matrix (see *SI Text*) and the *N*×*D* sample dataset and the runtime is therefore expected to be 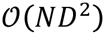.

### Datasets and Preprocessing

All data preprocessing was done in the R programming language using the *phyloseq* package for analysis of microbiome census data (80) as well as the the *ape* (81) and *phangorn* (82) packages for analysis of phylogenetic trees. For each of the three 16s rRNA datasets (consisting of an Operational Taxonomic Unit (OTU) table, taxonomic classifications, and phylogenetic tree) the preprocessing pipeline was as follows: 1) If unrooted, manually root the phylogenetic tree by specifying an outgroup, 2) resolve multichotomies, if present, with the function multi2di from the *ape* package which replaces multichotomies with a series of dichotomies with one (or several) branch(es) of length zero, 3) filter low abundance taxa and prune the tree accordingly, 4) filter low abundance samples, 5) add a pseudocount of 1 prior to PhILR transformation to avoid taking log-ratios with zero counts. The PhILR transform is robust to changing the value of this pseudocount (**Fig. S9**).

#### Human Microbiome Project (HMP)

This dataset was obtained from the QIIME Community Profiling Pipeline applied to high-quality reads from the v3-5 region, available at http://hmpdacc.org/HMQCP/. The phylogenetic tree was rooted with the phylum Euryarchaeota as an outgroup and multichotomies were resolved. Samples with fewer than 1000 counts were removed so that our analysis included the same samples as prior analyses (31). Taxa that were not seen with more than 3 counts in at least 1% of samples were removed. Samples from the left and right retroauricular crease and samples from the left and right antecubital fossa were grouped together, respectively, as preliminary PERMANOVA analysis suggested that these sites were indistinguishable (data not shown).

#### Global Patterns

The Global Patterns dataset was originally published in Caporaso, *et al.* (1). This dataset is provided with the *phyloseq* package and our preprocessing followed the methods outlined in McMurdie and Holmes (80). Specifically, taxa that were not seen with more than 3 counts in at least 20% of samples were removed, the sequencing depth of each sample was standardized to the abundance of the median sampling depth, and finally taxa with a coefficient of variation ≤ 3.0 were removed (80). The tree was rooted with Archaea as the outgroup and no multichotomies were present.

#### Costello Skin Sites (CSS)

The original dataset was collected by Costello et al. (30). A subset of the dataset containing only the skin samples was introduced as a benchmark for supervised machine learning by Knights et al. (34). This dataset was obtained from http://knightslab.org/data. The CSS dataset consists of counts from the v2 region of bacterial 16s rRNA genes. The tree was rooted at OTU 12871 (from phylum Plantomycetes) and multichotomies were resolved. This dataset had lower sequencing depth than the other two benchmarks. To retain a reasonable number of taxa while still removing potential spurious reads, we chose to filter taxa that were not seen with greater than 10 counts across the skin samples.

### Benchmarking

#### Distance/Dissimilarity Based Analysis

Distance between samples in PhILR transformed space was calculated using Euclidean distance. All other distance measures were calculated using *phyloseq* on the preprocessed data without adding a pseudocount. Principle coordinate analysis was performed for visualization using *phyloseq.* PERMANOVA was performed using the function *adonis* from the R package *vegan* (v2.3.4). Standard errors were calculated using bootstrap resampling with 100 samples each. Differences between the performance of Euclidean distance in PhILR transformed space and that of each other measure on a given task was tested using two-sided t-tests and multiple hypothesis testing was accounted for using FDR correction.

#### Supervised Classification

The performance of PhILR transformed data was compared against data preprocessed using one of two standard strategies for normalizing sequencing depth: the preprocessed data was transformed to relative abundances (*e.g*., each sample was normalized to a constant sum of 1; *raw*); or, a pseudocount of 1 was added, the data was transformed to relative abundances, and finally the relative abundances were log-transformed (*log)*.

All supervised learning was implemented in Python using the following libraries: *Scikit-learn* (v0.17.1), *numpy* (v1.11.0) and *pandas* (v0.17.1). Four classifiers were used: penalized logistic regression, support vector classification with RBF kernel, random forest classification, and k-nearest-neighbors classification. Each classification task was evaluated using the mean and variance of the test accuracy over 10 randomized test/train (30/70) splits which preserved the percentage of samples from each class at each split. For each classifier, for each split, the following parameters were set using cross-validation on the training set. Logistic regression and Support Vector Classification: the *‘C’* parameter was allowed to vary between 10^−3^ to 10^3^ and multi-class classification was handled with a one-vs-all loss. In addition, for logistic regression the penalty was allowed to be either *l*_1_ or *l*_2_. K-nearest-neighbors classification: the ‘weights’ argument was set to ‘distance’. Random forest classification: each forest contained 30 trees and the ‘max_features’ argument was allowed to vary between 0.1 and 1. All other parameters were set to default values. Due to the small size of the Global Patterns dataset, the supervised classification task was simplified to distinguishing human vs. non-human samples. Differences between each methods’ accuracy in a given task was tested using two-sided t-tests and multiple hypothesis testing was accounted for using FDR correction.

### Identifying balances that distinguish sites

To identify a sparse set of balances that distinguish sampling sites, we fit a multinomial regression model with a grouped l_1_ penalty using the R package *glmnet (v2.0.5*). The penalization term lambda was set by visually inspecting model outputs for clear bodysite separation (lambda=0.1198). This resulted in 35 balances with non-zero regression coefficients. Phylogenetic tree visualization was done using the R package *ggtree* (83).

### Variance and Depth

The orthonormality of PhILR balances and their association with internal nodes of the phylogenetic tree enabled us to investigate how the association between neighboring clades varied with phylogenetic depth. The original variance of log-ratios proposed by Aitchison as a measure of association (5) are not comparable on the same scale (it is unclear what constitutes a large or small variance) (4). However, the variance of ILR coordinates can be compared on the same scale because of the unit length of ILR basis elements (10).

We computed the variance of balances that did not include taxa weights (*i.e.*, **p** = (1,…,1)). We also omitted branch length weights 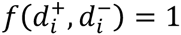 when computing balance variances to avoid directly weighting variances by phylogenetic distances. We omitted taxa weights because of concern that zero values would vary as a function of phylogenetic depth and could therefore systematically bias our analysis of balance variance as a function of phylogenetic depth. That is, balances closer to the root of the tree have more descendant tips that could be non-zero compared to balances closer to the tips of the tree. We therefore calculated balance values 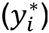 on non-zero counts. In practice, we retained balances that met the following criteria: the term 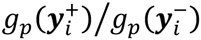 had non-zero counts from some part(s) within the subcomposition 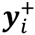 and some other part(s) within the subcomposition 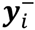 in at least 40 samples from that body site. To further focus our analysis of each HMP body site on non-zero counts, prior to calculation of balance values, taxa present in less than 20% of samples from that site were excluded and subsequently samples that had less than 50 total counts were excluded.

In order to investigate the overall relationship between balance variance and phylogenetic depth we used linear regression. A balance’s depth in the tree was calculated as its mean distance to its descendant tips (*d*). For a given body site the following model was fit:

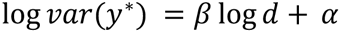

where *d* represents mean distance from a balance to its descendant tips. We then set out to test the null hypothesis that β = 0, or that the variance of the log-ratio between two clades was invariant to the distance of the two clades from their most recent common ancestor. For each site, a null distribution for β was constructed by permutations of the tip labels of phylogenetic tree. We chose this permutation scheme to ensure that the increasing variance we saw with increasing proximity of a balance to the root was not because deeper balances had more descendant tips, an artifact of variance scaling with mean abundance, or due to bias introduced due to our handling of zeros. Furthermore, the null distribution for β is symmetric about β = 0 which further supports that balance variance depends on phylogenetic depth through an ecological mechanism and not through a statistical artifact (**Fig. S10**). Two tailed p-values were calculated for β based on 20000 samples from each site’s respective null distribution. FDR correction was applied to account for multiple hypothesis testing between body sites.

To visualize local trends in the relationship between balance variance and phylogenetic depth, a LOESS regression was fit independently for each body site. This was done using the function *geom_smooth* from the R package ggplot2 (v2.1.0) with default parameters.

### Integrating Taxonomic Information

Taxonomy was assigned to OTUs in the HMP dataset using the *assign_taxonomy.py* script from *Qiime* (v1.9.1) to call *uclust* (v1.2.22) with default parameters using the representative OTU sequences obtained as described above. Taxonomic identifiers were assigned to the two descendant clades of a given balance separately using a simple voting scheme and combined into a single name for that balance. The voting scheme occurs as follows: (1) for a given clade, the entire taxonomy table was subset to only contain the OTUs that were present in that clade (2) starting at the finest taxonomic rank the subset taxonomy table was checked to see if any species identifier represented ≥95% of the table entries at that taxonomic rank, if so that identifier was taken as the taxonomic label for the clade (3) if no consensus identifier was found, the table was checked at the next most-specific taxonomic rank.

Median phylogenetic depths for each taxonomic rank was estimated by first decorating a phylogenetic tree with taxonomy information using *tax2tree* (v1.0) (84). For a given taxonomic rank the mean distance to tips was calculated for each internal node possessing a label that ended in that rank. The median of these distances was used to display an estimate of the phylogenetic depth of that given rank. This calculation of median phylogenetic depth of different taxonomic ranks was done separately for each body site.

## Acknowledgements

We thank Jesse Shapiro, Aspen Reese, Firas Midani, Heather Durand, Jonathan Friedman, Susan Holmes, and Simon Levin for their helpful comments, Dan Knights for providing us with the CSS dataset, and Klaus Schliep and Liam Revell for their insight into manipulation of phylogenetic trees in the R programming language. JS was supported in part by the Duke University Medical Scientist Training Program (GM007171). LAD was supported by the Global Probiotics Council, a Searle Scholars Award, and an Alfred P. Sloan Research Fellowship.

